# Quantification of bone loss, periosteal bone formation and novel histopathological changes in a mouse implant-related *Staphylococcus aureus* infection model

**DOI:** 10.64898/2026.08.05.742940

**Authors:** Qi Sun, Dzenita Muratovic, Helen Tsangari, Rebecca K. Sawyer, Mohammad A. Hossain, L. Bogdan Solomon, Paul H. Anderson, Gerald J. Atkins

**Affiliations:** Biomedical Orthopaedic Research Group, Discipline of Orthopaedics and Trauma, School of Medicine, College of Health, Adelaide University, Adelaide, SA, Australia; School of Pharmacy and Biomedical Sciences, College of Health, Adelaide University, SA, Australia; Department of Orthopaedics and Trauma, Royal Adelaide Hospital, Adelaide, SA, Australia; Discipline of Orthopaedics and Trauma, School of Medicine, College of Health, Adelaide University, Adelaide, SA, Australia

**Keywords:** implant-related infection, prosthetic joint infection, osteomyelitis, preclinical model, orthopaedic, osteocyte

## Abstract

Implant-associated bone infection involves a complex interplay between pathogenic stimuli and host cell responses, yet analysis in preclinical models has typically relied on qualitative or semi-quantitative measures. We aimed to establish a quantified evaluation framework to define host-pathogen relationships in a preclinical implant infection model. *Staphylococcus aureus-*coated stainless-steel implants were inserted trans-cortically in mouse tibiae and bone changes recorded longitudinally by *in vivo* micro-CT. An automated segmentation task list was developed to independently isolate and quantify cortical, periosteal-reactive, and trabecular bone compartments. RGB trichrome histomorphometry was used to quantify bone matrix integrity, osteocyte lacunar geometry, and osteoclastic activity. Droplet digital PCR was used to determine absolute bacterial and host genome copy number. Infected implants produced marked reductions in trabecular bone volume fraction, number, and bone mineral density (BMD), together with decreased cortical bone volume fraction and increased cortical porosity, accompanied by significant elevations in periosteal bone volume fraction. Histologically, infected bone exhibited increased eroded surface indicative of osteoclastic resorption, extensive degraded bone matrix and pathological remodelling of osteocyte lacunae towards circularity, consistent with an osteocytic osteolysis response. Infection-induced changes to cortical bone structure correlated mostly with host cell rather than bacterial load; however, cortical BMD negatively correlated with the bacterial:host genome ratio. This multifaceted, quantified framework reveals distinct pathobiological effects of implant-associated infection on trabecular, cortical, and periosteal bone compartments, bone matrix and osteocyte and osteoclast populations, consistent with reports in human patients, suggesting that major pathological changes are driven by the host bone cell response to infection.

## Introduction

Osteomyelitis in juveniles or adults, either spontaneous or following musculoskeletal trauma, surgery, or prosthesis implantation (periprosthetic joint infection; PJI), is characterised by both bone loss (osteolysis) and a sclerotic bone formation response [1]. In human PJI bone, osteolysis has been attributed to both the bone-resorbing activity of osteoclasts and a recently discovered bone matrix degradation response by osteocytes [2]. Our recent study further showed that specific alterations to the osteocyte lacunocanalicular network, in terms of increased osteocyte lacunar circularity and degradation of peri-lacunocanalicular bone collagen matrix, among others, were predictive of PJI compared to aseptic osteolysis and primary joint replacement cohorts [3]. Given the uncontrolled nature and general difficulty in obtaining clinical specimens, preclinical models are still considered the cornerstone for investigating the *in vivo* pathogenesis and potential therapeutic interventions of osteomyelitis [4–7]. A variety of species, including mice [8], rats [9], rabbits [10], pigs [11] and sheep [7] have been widely utilised in this field. The establishment of these models relies on two primary methodologies: direct bacterial inoculation of either the bone or the bone implant [8], or hematogenous seeding via intravenous injection [11].

The toolkit for evaluating the extent and progression of infection in preclinical models should ideally be multifaceted, including haematological analysis to monitor systemic inflammation, micro-computed tomography (micro-CT) imaging to assess architectural bone changes, and histopathological evaluation to characterise tissue-level changes. For micro-CT-based bone analysis, precise segmentation of cortical and trabecular bone is a prerequisite for robust data interpretation. Currently, both manual and various automated segmentation methods are employed. Although manual methods are historically reliable, with inter-analyser differences under 2% [12] and the discrepancy between manual and automated outputs having low bias [13], they are nevertheless open to user variation and are also time consuming. In contrast, automated segmentation offers substantial advantages by eliminating subjectivity, enhancing time efficiency, and providing a highly reproducible and scalable framework for evaluating complex bone structures [12, 14, 15].

Comprehensive histopathological evaluation of osteomyelitis integrates bacterial staining with the characterisation of localised inflammatory cell infiltration. Clinically and experimentally, histomorphometric quantification of polymorphonuclear neutrophils (PMNs) via standard haematoxylin and eosin (H&E) staining, or scoring systems assessing necrosis and granulocyte infiltration, remains a direct and essential determinant for defining active tissue-level infection [16]. For pathogen assessment, a multi-staining approach, including Gram-staining, Periodic Acid-Schiff, and Grocott’s Methenamine Silver, alongside specific antibody immunochemistry, has been employed to visualise bacterial distribution but not to quantify bacterial load [17]. However, while these established scoring matrices provide exceptional diagnostic power regarding the host immune/inflammatory response, they offer limited direct insight into the concurrent degradation of the bone matrix and the involvement of the resident bone cells. Mounting evidence demonstrates that osteocytes, the most abundant cells in bone, are not passive bystanders but rather active participants in the pathogenesis of bone infection. *S. aureus* has been shown to directly invade and colonise the osteocyte lacunocanalicular network (OLCN) [18, 19], triggering pathological host cellular adaptations and inducing osteocytic osteolysis, a process characterised by the active degradation of peri-lacunocanalicular collagen matrix and substantial lacunar geometric remodelling [2, 3]. Consequently, capturing these microenvironmental changes necessitates a quantitative histological framework for evaluating matrix integrity alongside osteocyte morphology to complement traditional inflammatory scoring.

To bridge the gap and provide a multi-angled evaluation of implant-associated osteomyelitis, this study introduces an integrated multi-scale framework using a mouse model. Distinct from conventional qualitative methods, we employed automated micro-CT segmentation to capture cortical and trabecular bone dynamics. Furthermore, we validated the utility of RGB trichrome staining for the direct quantitative analysis of bone matrix degradation and the osteocyte lacunar morphology for the first time in non-human bone. Finally, droplet digital PCR (ddPCR) was implemented for the absolute quantification of both bacterial and host cell load, and the bacteria: host genome ratio present within tissue sections [18]. This multi-angle approach seeks to define the bone structural and histological changes that occur in implant-related infection and elucidate whether architectural bone destruction is primarily dictated by the pathogen load or synchronised with the host intrinsic cellular responses.

## Materials and Methods

### Staphylococcus aureus preparation and mouse model

A trans-tibial implant-associated infection model in healthy 8-week male and female C57BL/6 mice was adapted from the method by Dan Li *et al.* [8]. The overall workflow used is depicted in **Figure 1**. Bacterial culture, stainless-steel pin (Australian Entomological Supplies, South Murwillumbah, NSW, Australia) coating and surgical procedure were performed as previously described [18]. The *S. aureus* reference strain ATCC25923 was used in this study as it previously exhibited the most reproducible implant attachment kinetics [18]. In brief, a single colony was inoculated into broth and cultured to reach the 1×10^9^ CFU/ml plateau phase. Sterile stainless-steel pins with a 0.3 mm diameter and 38 mm length were immersed in the bacterial suspension for 20 min to allow for surface attachment. Pins coated with sterile saline were used as implants in control mice. Mice were randomly assigned to either the infection group (*S. aureus*-coated pins, n = 12, 6 males and 6 females) or the control group (sterile saline-immersed pins, n = 12, 6 males and 6 females). Analgesia was administered perioperatively, and surgical sites (left tibia) were shaved and disinfected with Betadine before implantation. *S. aureus* or sterile saline-coated pins were inserted trans-cortically with a length of 6 mm of pin remaining in the bone. Post-surgical pain relief (0.3 mg/ml Buprenorphine) was administered subcutaneously for 2-4 hours, depending on clinical signs of pain. Animal health and wellbeing were monitored daily throughout the experimental period. The use of mice and all associated procedures were approved by the Institutional Animal Research Ethics Committee (Approval No. U20-23).

**Figure 1.**
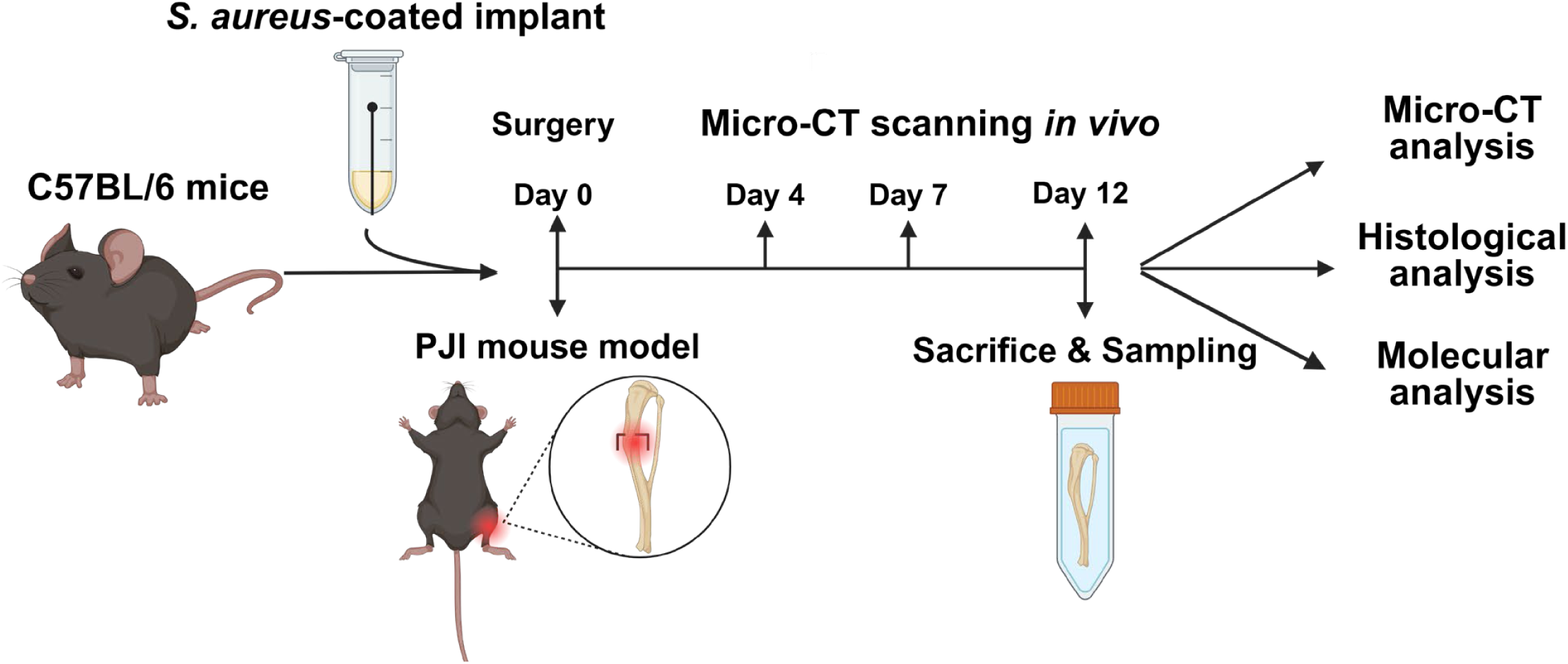
Experimental workflow of the PJI mouse model. (A) Schematic workflow of the experimental design. C57BL/6 mice underwent surgery with the trans-tibial implantation of *S. aureus*-coated implant to establish the PJI model. *In vivo* micro-CT scanning was performed on day 4, 7, and 12 post-surgery, followed by sacrifice for histological and molecular analyses.

### Image acquisition and definition of the volume of interest (VOI)

To longitudinally evaluate operated tibia bone changes, *in vivo* micro-CT (Skyscan 1276, Bruker) was performed on post-surgery days 4, 7, and 12. Scans were conducted at 70 KV and 200 μA with an isotropic voxel size of 8 μm. To ensure quantitative accuracy and mitigate potential scanner drift, density calibration was performed prior to scanning using two standard Phantoms containing known concentrations of Calcium Hydroxyapatite (CaHA; 0.25 and 0.75 g/cm^3^). The 3D datasets from different time points were spatially aligned using DataViewer software (Bruker) to ensure consistent anatomical positioning and fit across all longitudinal scans. To ensure that the same anatomical region was analysed across all time points, a standardised volume of interest (VOI) selection strategy based on anatomical landmarks was implemented (**Fig. 2A**). The proximal growth plate of the tibia and the superior margin of the implant pinhole were defined as the upper and lower reference landmarks, respectively. For the day 4 baseline analysis, the VOI was defined as a 60-slice region starting at an offset of 10 slices proximal to the superior margin of the pinhole. For subsequent time points (day 7 and 12), the analysis region was dynamically mapped by calculating the relative anatomical distance between the growth plate and the pinhole margin as established on day 4. This approach ensured that the VOI remained consistent with the initial target bone tissue throughout the 12 day experimental period.

**Figure 2.**
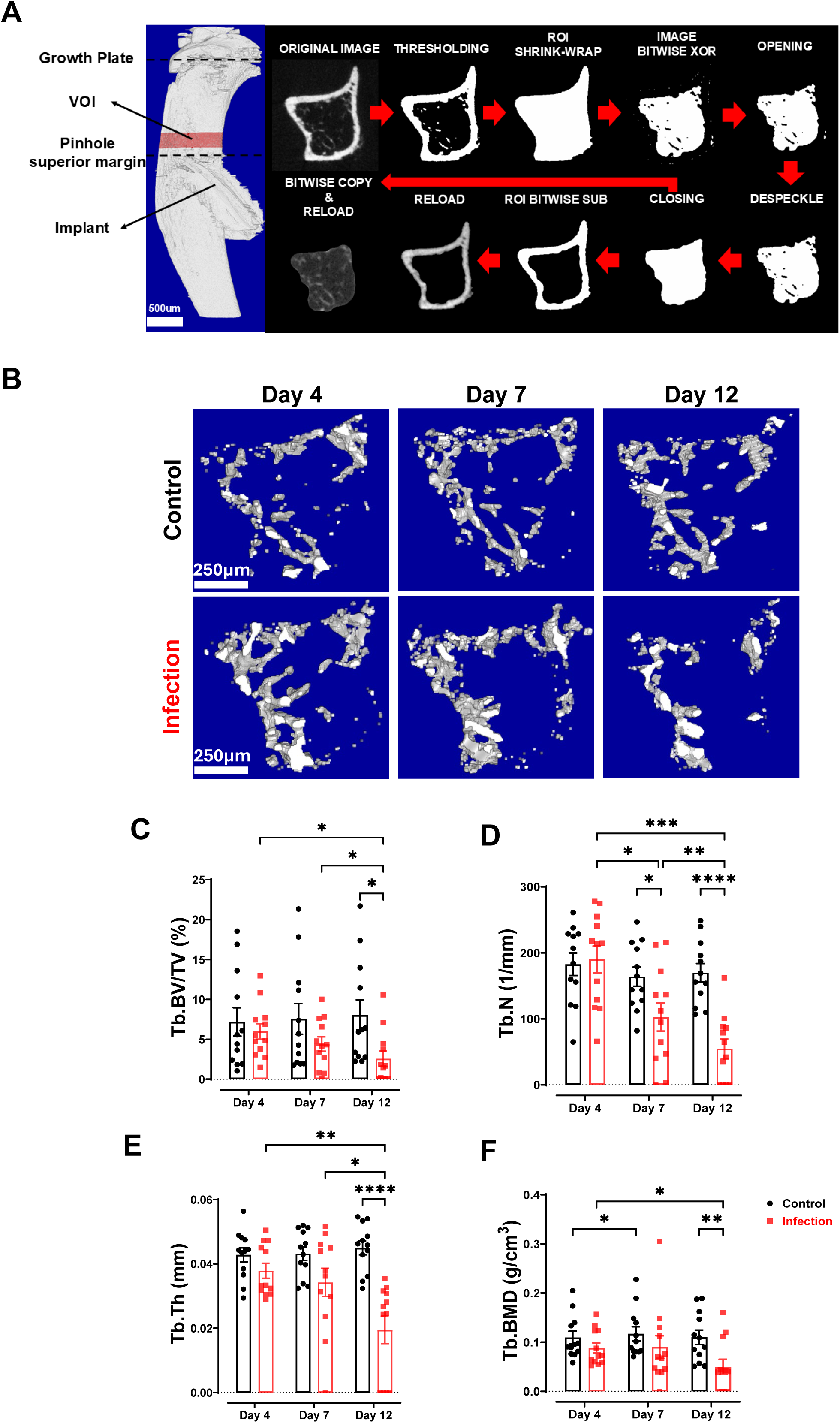
Micro-CT VOI selection, automated segmentation and analysis strategy. (A) Schematic of the tibial VOI definition based on anatomical landmarks. The flow diagram illustrates the task-list-based separation of cortical and trabecular bone. (B) Representative 3D micro-CT reconstructions of the trabecular bone within the defined ROI above the surgical site. Images show longitudinal changes in bone volume and architecture in the control and infection groups at days 4, 7, and 12. (C) Quantitative micro-CT analysis of trabecular bone parameters, including Tb.BV/TV (%), Tb.Th (mm), Tb.N (1/mm), and Tb.BMD (g/cm^3^). Data are presented as mean ± SEM (n=12 mice per group, pooled from 6 males and 6 females). Statistical significance was determined by Two-way ANOVA followed by Tukey’s multiple comparisons test. \**p*<0.05, \*\**p*<0.01, \*\*\**p*<0.001, \*\*\*\**p*<0.0001 indicates a significant difference between the indicated groups.

### Automated segmentation and morphometric analysis

To eliminate user-dependent bias and ensure high reproducibility, a standardised, automated segmentation task-list was developed within CTan software (Bruker, Belgium) to independently isolate the trabecular (Tb), original cortical (Ct) and combined cortical and periosteal reactive (Ct.Ps) bone fractions. Initially, optimal binarisation thresholds for each bone compartment were determined by visual inspection and histogram analysis using the binary function in CTan, ensuring accurate coverage of mineralised tissue while minimising artifact. Following this calibration, a Gaussian filter was applied to the raw reconstructed datasets to minimise high-frequency noise, and mineralised tissues were binarised via global thresholding with attenuation ranges tailored to each compartment: 85–255 for Tb, 160–255 for Ct and 125–255 for Ct.Ps. The isolation of the cortical compartments was achieved through a precise sequence of morphological bit-map operations. A region of interest (ROI) ‘shrink-wrap’ operation was first performed to enclose the entire cross-sectional area, followed by a bitwise Exclusive OR (XOR) operation (Image = ROI XOR Image) to isolate the internal endocortical space. To refine this internal region, morphological Opening (Round kernel) and a despeckling operation were applied to remove non-contiguous noise, followed by a morphological Closing (Round kernel) to fill all internal black pixels, thereby creating a solid internal mask. The final cortical region (Ct or Ct.Ps) was then extracted by a bitwise Subtract (SUB) operation (ROI = ROI SUB Image). For the trabecular compartment, the region was generated using a COPY image operation based on the previously defined endocortical boundaries to ensure anatomical alignment with the cortical masks (**Fig. 2A**).

Following the automated compartmentalisation, the designated bone masks were reloaded for high-resolution 3D analysis. To capture fine architectural details, an adaptive thresholding approach was implemented with specific grey-level ranges: 125–255 for Ct.Ps, 160–255 for Ct, and 85–255 for Tb. Quantitative 3D morphometric parameters were then calculated based on the binarised datasets according to standard nomenclature. The processed bitmaps were exported for volumetric assessment and subsequent 3D visualisation. Representative 3D reconstructions were generated using CT Vox software (Bruker) to provide a high-fidelity representation of the infection-induced osteolysis and reactive periosteal bone formation across the longitudinal time points.

### Bone specimen processing, RGB staining and analysis

At 12 days post-operation, mice were deeply anesthetised and humanely euthanised by cervical dislocation. The operated tibias were harvested and immediately fixed and decalcified in TheraLin fixative (Grace Biolabs, Bend, OR, USA) at room temperature for 7 days. Decalcified tibias were then processed for routine paraffin embedding and sectioning.

Tibial sections were stained using RGB trichrome staining, as previously described [3, 20] with some modifications to suit mouse bone. Sections were dewaxed and dehydrated, followed by sequential staining with 1% Alician Blue in 3% aqueous acetic acid (pH 2.5) for 20 minutes, 0.04% Fast Green for 20 minutes, and 0.1% Sirius Red in aqueous Picric acid for 1.5 minutes with differentiation in 1% acetic acid. All stained sections were imaged using a NanoZoomer scanner (Hamamatsu Photonics, Shizuoka, Japan), and RGB images were processed in Fiji software (open source, Java-based). Bone matrix degradation and osteocyte lacunae morphology were analysed, as previously described [3]. For bone matrix degradation, RGB-stained sections were imaged and processed using colour deconvolution to separate stain components. Degraded bone matrix (%DBM) was quantified as the proportion of degraded collagen (*red* stain), expressed as a percentage of total area. For characterisation of the osteocyte lacunae, image segmentation was performed using the Trainable Weke Segmentation plugin in Fiji, which utilises supervised machine learning algorithms to classify objects based on pixel features. The classifier was specifically trained to distinguish between osteocyte lacunae (Class 1) and bone matrix background (Class 2). Separate binary masks were generated for the lacunar compartment, which were subjected to morphometric analysis.

### DNA template preparation and ddPCR analysis

Single 20 μm sections of the same embedded left operated tibial specimens used for histological analysis above were placed in microcentrifuge tubes and deparaffinised using xylene and graded ethanol washes. The tissue was then subjected to DNA isolation by the addition of 200µl DirectPCR^TM^ buffer (Viagen Biotech) containing 200 µg/ml proteinase K (ThermoFisher Scientific) at 55℃ overnight, followed by heat-inactivation at 85℃ for 15 min to terminate enzymatic digestion. The quantification of host bone cell and bacterial genome copies was performed using a QX200 ddPCR system, according to the manufacturer’s instructions and as previously described [18, 21]. Primers targeting a mouse genome-specific sequence within the single-copy type X collagen gene (*Col10a1*) and *S. aureus* genome sequence within the single copy *tuf* gene were used for the amplification of host and bacterial signals, respectively. The sequences of all primer sets used are listed in **Supplementary Table 1**.

### Statistical analysis

Data are presented as means ± standard error of the mean (SEM), unless stated otherwise. Statistical analyses were performed using GraphPad Prism 10.2 (GraphPad Software, San Diego, CA, USA). Longitudinal micro-CT parameters were evaluated using a two-way ANOVA to compare differences between groups and across time-points. Quantitative histological data from RGB staining were analysed on day 12 specimens using two-way ANOVA or multiple comparisons with Tukey’s correction. For the absolute quantification of bacterial and host genome copy numbers, and the bacterial:host genome ratio, differences between groups were assessed using a one-way ANOVA. For correlation analysis, to determine the relationship between bacterial load and infection-induced bone alterations, Pearson correlation coefficients (*r*) were calculated. Linear regression and correlation matrices were utilised to evaluate the association between measures, including micro-CT morphometric indices and histological eroded surface (ES/BS%), %DBM and osteocyte lacunar circularity (Lac.Circ). In all analyses, a value for *p* < 0.05 was considered statistically significant.

## Results

### Behaviour of the mouse model

To investigate the progression of bone destruction, a localised bone implant-associated infection model was utilised in C57BL/6 mice using *S. aureus*-coated trans-tibial implants, with multi-modal assessment performed on postoperative days 4, 7 and 12 (see **Fig. 1**). Control mice received sterile saline-immersed implants as uninfected controls. Body weight and temperature were monitored to evaluate overall host physiological stability in response to the procedure and infection. Both control and infection groups exhibited a consistent pattern of body weight maintenance and steady growth over the 12-day experimental period across both male and female cohorts (**Suppl. Fig. 1**). Additionally, body temperature in the infection group remained within the normal physiological range after transient postoperative fluctuations (**Suppl. Fig. 1B**).

### Longitudinal in vivo micro-CT evaluation of trabecular bone

Quantitative 3D analysis revealed progressive deterioration of the trabecular bone architecture in the infection group compared to the control group (**Fig. 2B**). While there were no significant differences in trabecular parameters at day 4, progressive structural loss became evident over time in the infection group (**Fig. 2C-F)**. By day 12, the infection group exhibited significant reductions in trabecular bone volume fraction (Tb.BV/TV, *p* < 0.05), trabecular thickness (Tb.Th, *p* < 0.0001), and trabecular bone mineral density (Tb.BMD, *p* < 0.01) compared to time-matched controls (**Fig. 2C, E, F)**. Most prominently, trabecular number (Tb.N) in infected mice was decreased at days 7 and 12 relative to controls (**Fig. 2D**). Separate subgroup analyses in male (**Suppl. Fig. 2A**) and female (**Suppl. Fig. 3A**) cohorts demonstrated consistent longitudinal trends in trabecular bone loss, confirming that these architectural alterations occur similarly across sexes.

### Impact of infection on cortical bone structural integrity

Representative 3D reconstruction and micro-CT analysis revealed progressive pathological alterations in the cortical bone compartment following infection (**Fig. 3**). Visual inspection of the segmented Ct showed increasing surface irregularities and localised destruction, including endosteal and periosteal erosion, as well as increased cortical porosity, in the infection group from days 4 to 12 (**Fig. 3A**, top row). Substantial periosteal reactive bone formation in the infection group was evident at days 7 and 12 (**Fig. 3A**, middle row). Merged visualisations (**Fig. 3A**, bottom row) further delineated the spatial relationship between the underlying cortical porosity and the overlying reactive periosteal bone.

**Figure 3.**
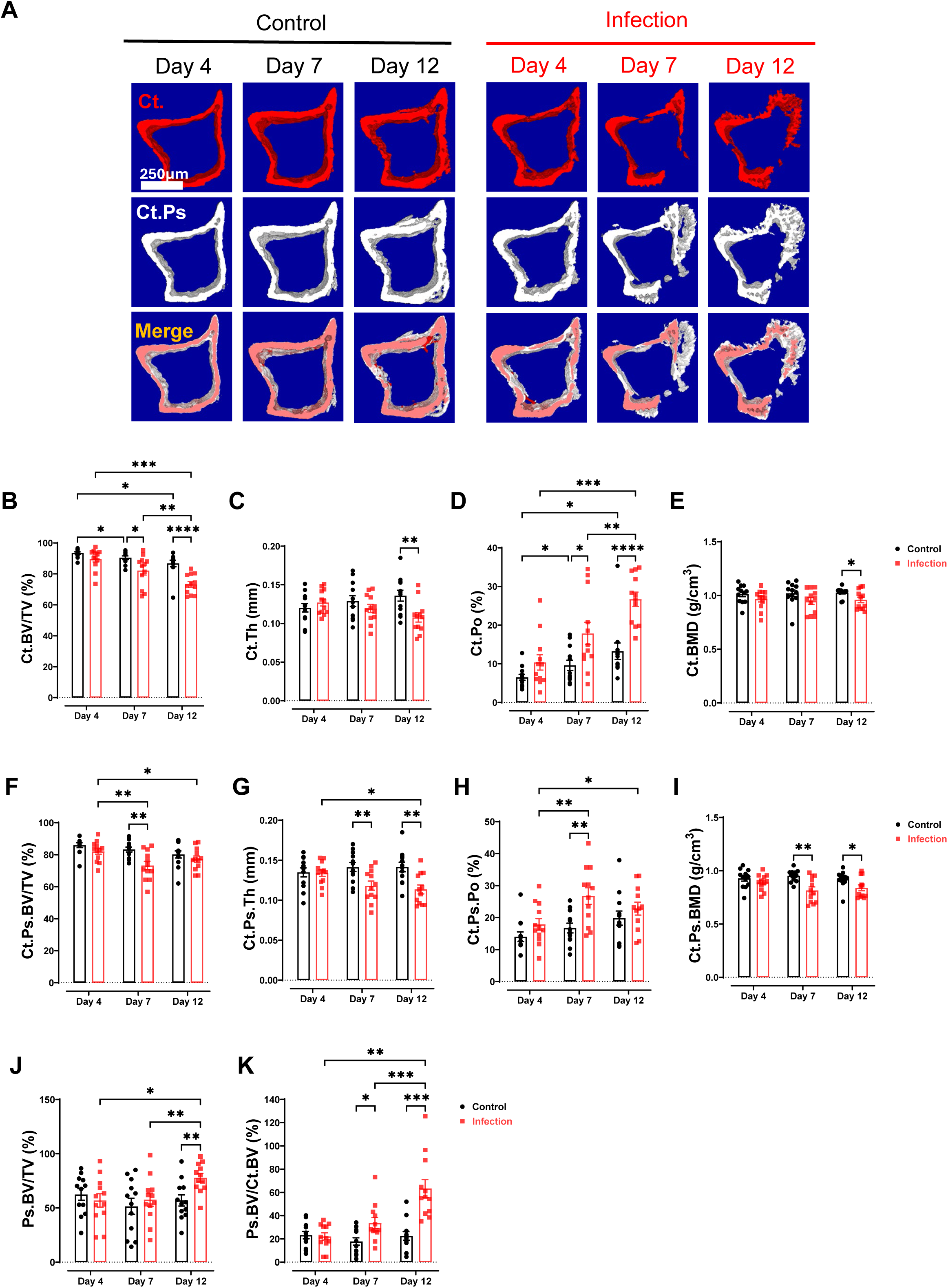
Micro-CT evaluation of cortical bone changes and periosteal bone formation. (A) Representative micro-CT illustrating the dynamic alterations in Ct and Ps within the same VOI as in Figure 2A at day 4, 7, and 12 post-surgery. The top row (Ct, red) displays the cortical bone excluding the periosteal component; the middle row (Ct.Ps, grey) shows the cortical bone inclusive of periosteal reactive bone; the bottom row (Merge) provides a superimposed visualisation to facilitate comparison between the original cortex and the newly formed bone. (B) quantitative analysis of the cortical bone excluding periosteal bone, characterised by Ct. BV/TV (%), Ct.Th (mm), Ct.Po (%), and Ct.BMD (g/cm^3^). (C) Quantitative analysis of Ct.Ps.BV/TV (%), Ct.Ps.Th (mm), Ct.Ps.Po (%), and Ct.Ps.BMD (g/cm^3^). (D) Assessment of periosteal bone formation, showing the Ps.BV/TV (%) and Ps.BV/Ct.BV (%). Data are expressed as mean ± SEM (n=12 mice per group, pooled from 6 males and 6 females). Statistical significance was determined by Two-way ANOVA followed by Tukey’s multiple comparisons. \**p*<0.05, \*\**p*<0.01, \*\*\**p*<0.001, \*\*\*\**p*<0.0001 indicate significant differences between the control and infection groups at specific time points, or between different time points within the same group.

Quantitative analysis of Ct compartments confirmed progressive osteo-morphological alterations following infection. Compared to time-matched controls, the infection group exhibited statistically significant reductions in cortical bone volume fraction (Ct.BV/TV) starting at day 7 and worsening by day 12 (*p* < 0.0001; **Fig. 3B**). By day 12, cortical thickness (Ct.Th) in infected mice was also reduced compared to controls (*p* < 0.01; **Fig. 3C**). Correspondingly, cortical porosity (Ct.Po) exhibited a marked elevation in infected mice at both day 7 (*p* < 0.05) and day 12 (*p* < 0.0001) relative to control mice (**Fig. 3D**). Cortical bone mineral density (Ct.BMD) was significantly decreased at day 12 (*p* < 0.05; **Fig. 3E**).

Quantitative micro-CT analysis of the Ct.Ps compartment captured significant transient and sustained pathological shifts. In infected mice, Ct.Ps.BV/TV significantly dropped at day 7 compared to controls (*p* < 0.01; **Fig. 3F**). Ct.Ps thickness (Ct.Ps.Th) was significantly reduced in the infection group at both day 7 (*p* < 0.01) and day 12 (*p* < 0.01) relative to control mice (**Fig. 3G**). Furthermore, Ct.Ps porosity (Ct.Ps.Po) demonstrated a significant elevation at day 7 (*p* < 0.01; **Fig. 3H**), while Ct.Ps bone mineral density (Ct.Ps.BMD) was significantly lower in infected mice at day 7 (*p* < 0.01) and day 12 (*p* < 0.05) compared to controls (**Fig. 3I**).

Evaluation of the periosteal bone formation (Ps) compartment revealed robust reactive bone expansion in response to infection. By day 12, the infection group exhibited a significant increase in periosteal bone volume fraction (Ps.BV/TV) compared to controls (*p* < 0.01; **Fig. 3J**). The ratio of periosteal bone volume to underlying cortical bone volume (Ps.BV/Ct.BV) in infected mice was significantly elevated as early as day 7 (*p* < 0.05) and was further elevated by day 12 (*p* < 0.001) compared to time-matched controls (**Fig. 3K**). Sex-specific subgroup analysis (**Suppl. Fig. 2B–D** for males; **Suppl. Fig. 3B–D** for females) mirrored these cortical and periosteal structural dynamics.

### RGB trichrome staining assessment of bone matrix degradation and osteocyte lacunar morphology

To evaluate the impact of implant-associated bone infection with *S. aureus* on the bone matrix and cellular microenvironment, RGB trichrome staining was performed on both the ipsilateral (operated, left; Ipsi) and contralateral intact (unoperated, right; Contra) tibiae in the control (CONT) and infected (INF) groups (**Fig. 4**, **Table 1**). Qualitative histological assessment of the ipsilateral tibia revealed marked degraded bone matrix in the infected group, indicated by predominantly red-stained degraded collagen over blue/green-stained mature collagen compared to the uninfected control group (**Fig. 4A**). Quantitative analysis confirmed that the %DBM in the infected group ipsilateral tibia [INF (Ipsi)] was significantly elevated compared to the corresponding control group [CONT (Ipsi), *p* < 0.0001, **Fig. 4B**], and the contralateral intact tibias of the infected group [INF (Contra), *p* < 0.0001, **Table 1**]. No significant difference in %DBM was found between the ipsilateral and contralateral tibia within the control group [CONT (Ipsi) vs CONT (Contra), *p* = 0.3992, **Table 1**].

**Figure 4.**
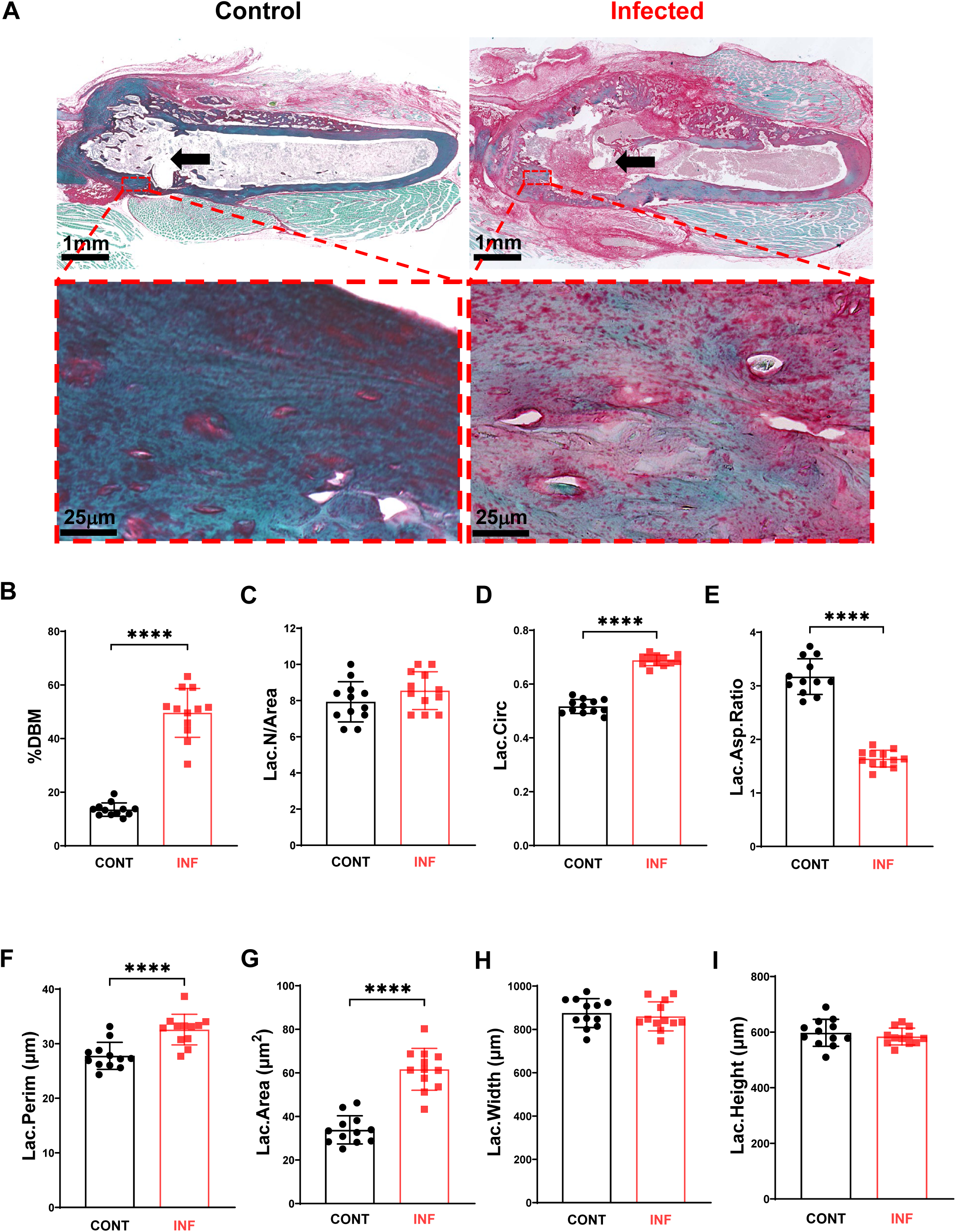
Histological and molecular evaluation of bone matrix alterations, lacunar morphology. (A) Representative RGB-staining images of the peri-implant bone in the control and infection groups. Red staining highlights degraded collagen, while blue/green staining indicates mature collagen. High-magnification images illustrate the morphological alterations of osteocyte lacunae within the bone matrix. Arrowheads indicate the implant hole. (B-I) Quantitative histological analysis of five selected regions of interest (ROI) surrounding the implant. The percentage of degraded bone matrix (DBM%), Lac.Circ, Lac.Asp.Ratio, Lac.Area, Lac.Perim, Lac.N/Area, Lac.Width and Lac.Height were evaluated across five ROIs, with no fewer than five lacunae analysed per region. Data are expressed as mean ± SEM. Statistical significance was determined by Two-way ANOVA followed by Tukey’s multiple comparisons. The ES/BS% was determined within the peri-implant bone area under 10× magnification. Data are presented as mean ± SEM, with statistical significance between individual mice determined by one-way ANOVA followed by multiple comparisons. \**p*<0.05, \*\**p*<0.01, \*\*\**p*<0.001, and \*\*\*\**p*<0.0001.

**Table 1.** Quantitative comparison of bone matrix degradation and osteocyte lacunocanalicular morphology between ipsilateral and contralateral tibiae in control and infection groups.

| **Group Comparison** | **Mean**<br>**1** | **Mean**<br>**2** | **Mean Difference (95%**<br>**CIs)** | **p-Value**<br>**(Tukey**<br>**Adjusted)** |
| --- | --- | --- | --- | --- |
| **% Bone matrix degradation** |  |  |  |  |
| CONT (Ipsi) – CONT (Contra) | 13.51 | 11.73 | 1.771 (-1.156 to 4.698) | 0.3992 |
| INF (Ipsi) – INF (Contra) | 49.58 | 12.38 | 37.20 (34.27 to 40.13) | <0.0001 |
| CONT (Ipsi) – INF (Ipsi) | 13.51 | 49.58 | -36.08 (-39.00 to -33.15) | <0.0001 |
| CONT (Contra) – INF (Contra) | 11.73 | 12.38 | -0.648 (-3.575 to 2.279) | 0.9397 |
| **Lacunar circularity** |  |  |  |  |
| CONT (Ipsi) – CONT (Contra) | 0.516 | 0.585 | -0.069 (-0.093 to 0.045) | <0.0001 |
| INF (Ipsi) – INF (Contra) | 0.687 | 0.646 | 0.041 (0.017 to 0.064) | <0.0001 |
| CONT (Ipsi) – INF (Ipsi) | 0.516 | 0.687 | -0.171 (-0.195 to -0.147) | <0.0001 |
| CONT (Contra) – INF (Contra) | 0.585 | 0.646 | -0.061 (-0.084 to -0.037) | <0.0001 |
| **Lacunar aspect ratio** |  |  |  |  |
| CONT (Ipsi) – CONT (Contra) | 3.174 | 2.957 | 0.217 (0.005to 0.429) | 0.0423 |
| INF (Ipsi) – INF (Contra) | 1.641 | 2.480 | -0.838 (-1.051 to -0.626) | <0.0001 |
| CONT (Ipsi) – INF (Ipsi) | 3.174 | 1.641 | 1.533 (1.321 to 1.745) | <0.0001 |
| CONT (Contra) – INF (Contra) | 2.957 | 2.480 | 0.477 (0.265 to 0.689) | <0.0001 |
| **Lacunar number per area** |  |  |  |  |
| CONT (Ipsi) – CONT (Contra) | 7.933 | 7.600 | 0.333(-0.698 to 1.365) | 0.8366 |
| INF (Ipsi) – INF (Contra) | 8.550 | 8.133 | 0.417(-0.615 to 1.448) | 0.7222 |
| CONT (Ipsi) – INF (Ipsi) | 7.933 | 8.550 | -0.617(-1.648 to 0.415) | 0.4104 |
| CONT (Contra) – INF (Contra) | 7.600 | 8.133 | -0.533(-1.565 to 0.498) | 0.5388 |
| ***Lacunar perimeter*** |  |  |  |  |
| CONT (Ipsi) – CONT (Contra) | 27.77 | 28.10 | -0.3398 (-2.287 to 1.607) | 0.9691 |
| INF (Ipsi) – INF (Contra) | 32.57 | 30.60 | 1.968 (0.0207to 3.1914) | 0.0465 |
| CONT (Ipsi) – INF (Ipsi) | 27.77 | 32.57 | -4.807 (-6.753 to -2.860) | <0.0001 |
| CONT (Contra) – INF (Contra) | 28.10 | 30.60 | -2.499 (-4.446 to -0.5522) | 0.0058 |
| ***Lacunar area*** |  |  |  |  |
| CONT (Ipsi) – CONT (Contra) | 33.83 | 38.74 | -4.912 (-11.42 to 1.595) | 0.2083 |
| INF (Ipsi) – INF (Contra) | 61.63 | 49.66 | 11.96 (5.455 to 18.47) | <0.0001 |
| CONT (Ipsi) – INF (Ipsi) | 33.83 | 61.63 | -27.80 (-34.31 to -21.29) | 0.0001 |
| CONT (Contra) – INF (Contra) | 38.74 | 49.66 | -10.93 (-17.43 to -4.419) | 0.0277 |
| ***Lacunar width*** |  |  |  |  |
| CONT (Ipsi) – CONT (Contra) | 875.4 | 688.7 | 186.7 (112.2 to 261.3) | <0.0001 |
| INF (Ipsi) – INF (Contra) | 860.3 | 680.2 | 180.1 (105.5 to 254.6) | <0.0001 |
| CONT (Ipsi) – INF (Ipsi) | 857.4 | 860.3 | 15.15 (-59.40 to 89.69) | 0.9526 |
| CONT (Contra) – INF (Contra) | 688.7 | 680.2 | 8.541 (-66.01 to 83.09) | 0.9909 |
| ***Lacunar height*** |  |  |  |  |
| CONT (Ipsi) – CONT (Contra) | 597.6 | 421.7 | 175.9 (120.3 to 231.4) | <0.0001 |
| INF (Ipsi) – INF (Contra) | 584.1 | 476.1 | 108.0 (52.45 to 163.5) | <0.0001 |
| CONT (Ipsi) – INF (Ipsi) | 597.6 | 584.1 | 13.56 (-41.98 to 69.11) | 0.9213 |
| CONT (Contra) – INF (Contra) | 421.7 | 476.1 | -54.33 (-109.9 to 1.213) | 0.0578 |
| ***Eroded surface %*** |  |  |  |  |
| CONT (Ipsi) – CONT (Contra) | 13.85 | 11.44 | 2.402 (-7.530 to 12.33) | 0.9129 |
| INF (Ipsi) – INF (Contra) | 41.11 | 13.60 | 27.51 (17.58 to 37.44) | <0.0001 |
| CONT (Ipsi) – INF (Ipsi) | 13.85 | 41.11 | -27.27 (-35.38 to -19.16) | <0.0001 |
| CONT (Contra) – INF (Contra) | 11.44 | 13.60 | -2.156 (-13.56 to 9.312) | 0.9563 |

Note: Data are presented as mean values for each group alongside Mean Difference with 95% Confidence Intervals (95% CIs) ( $n = 12$ mice per group, 6 males and 6 females). CONT, uninfected control group receiving sterile implants; INF, infection group receiving *S. aureus*-coated implants; Ipsi, ipsilateral (operated, left) tibia; Contra, contralateral (unoperated, right) tibia. Statistical comparisons were performed using Two-way ANOVA with Tukey's multiple comparisons test. Adjusted values for $p < 0.05$ were considered statistically significant.

In addition, the morphology of osteocyte lacunae was qualitatively altered in response to the presence of an infected implant (**Fig. 4A**). Quantitative morphometric analysis demonstrated that the infection group exhibited distinct pathological remodelling of the OLCN. While the number of osteocytes per bone area (Lac.N/Area) was not altered (**Fig. 4C**), the INF (Ipsi) group displayed a significant increase in lacunar circularity (Lac.Circ; *p* < 0.0001, **Fig. 4D**) accompanied by a concomitant decrease in the lacunar aspect ratio (Lac.Asp.Ratio; *p* < 0.0001, **Fig. 4E**) relative to CONT (Ipsi). Furthermore, infection induced lacunar enlargement, evidenced by significant increases in both lacunar perimeter (Lac.Perim; *p* < 0.0001, **Fig. 4F**) and lacunar area (Lac.Area; *p* < 0.0001, **Fig. 4G**) in INF (Ipsi) compared to CONT (Ipsi). In contrast, no statistically significant differences were evident between the INF (Ipsi) and CONT (Ipsi) groups regarding lacunar width (Lac.Width; *p* = 0.9526, **Fig. 4H**), or lacunar height (Lac.Height; *p* = 0.9213, **Fig. 4I**, **Table 1**). These results suggest that infection induces both significant and rapid bone collagen matrix breakdown and pathological remodelling of osteocyte lacunar geometry, consistent with reported findings in human PJI bone [3].

Intra- and inter-group comparisons involving the contralateral right tibia revealed subtle but distinct systemic effects of infection (**Table 1**). Within the infection group, the infected ipsilateral tibia exhibited significantly greater lacunar distortion than its contralateral counterpart [INF (Ipsi) vs INF (Contra)], as shown by increased Lac.Area (*p* < 0.0001), increased Lac.Circ (*p* < 0.0001) and decreased Lac.Asp.Ratio (*p* < 0.0001, **Table 1**). However, comparing the infection group’s contralateral tibia [INF (Contra)] with the control group’s contralateral tibia [CONT (Contra)], the unoperated tibia of infected mice exhibited significant increases in Lac.Area (*p* = 0.0277), Lac.Perim (*p* = 0.0058), and Lac.Circ (*p* < 0.0001), alongside a decreased Lac.Asp.Ratio (*p* < 0.0001, **Table 1**), supporting systemic osteocyte microenvironment alterations driven by an ostensibly localised distal infection.

Osteoclastic bone resorption activity was assessed in the same RGB-stained sections by histomorphometric measurement of eroded surface per bone surface (ES/BS%). The INF (Ipsi) group exhibited significantly higher osteoclastic bone resorption compared to both CONT (Ipsi) (*p* < 0.0001) and INF (Contra) (*p* < 0.0001, **Table 1**). No significant differences in ES/BS% were detected between the ipsilateral and contralateral tibiae of the control group [CONT (Ipsi) vs CONT (Contra), *p* = 0.9129], nor between the contralateral tibia of the control and infected groups [CONT (Contra) vs INF (Contra), *p* = 0.9563, **Table 1**].

### Molecular quantification of bacterial load in tissue sections via ddPCR

The absolute copy number of *S. aureus* genomes relative to mouse host cell genomes present within bone tissue sections from mice receiving an infected implant was measured using ddPCR. Absolute *S. aureus* genome copies were quantified across three separate tissue section levels per mouse, displaying localised intra-sample variability within individual animals while maintaining consistent bacterial burdens across the cohort (**Suppl. Fig. 4A**). Regarding host genome copies, variations were observed among individual mice, reflecting differences in total host tissue area across histological sections (**Suppl. Fig. 4B**). When normalised to host cellularity, the bacterial-to-host genome copy ratio showed no statistically significant differences among individual infected mice across sexes, demonstrating a stable relative microbial burden within the infected tissue microenvironment (**Suppl. Fig. 4C**). Baseline host genome copy numbers were similarly quantified in non-infected control mice (n = 12, comprising 6 males and 6 females) to establish physiological cellular baseline ranges (**Suppl. Fig. 4D**). No *S. aureus* genome signal was detected in control samples (data not shown).

### Baseline multi-parameter correlation analysis of bone matrix integrity, microarchitecture and host genome copy in the control group

To establish baseline physiological coupling under homeostatic conditions, multi-parameter Pearson correlation analyses were conducted across the trabecular (**Suppl. Fig. 5**) and cortical (**Suppl. Fig. 6**) compartments in the control group receiving sterile implants.

Within the trabecular compartment under physiological homeostasis (**Suppl. Fig. 5A**), Tb.BV/TV_CONT_ demonstrated significant positive linear correlation with Tb.Th_CONT_ (*r* = 0.645, *p* = 0.023) (**Suppl. Fig. 5B**), Tb.BMD_CONT_ (*r* = 0.885, *p* = 0.0001) (**Suppl. Fig. 5C**), and ES/BS%_CONT_ (*r* = 0.679, *p* = 0.015) (**Suppl. Fig. 5D**). Furthermore, Tb.Th_CONT_ positively correlated with Tb.BMD_CONT_ (*r* = 0.682, *p* = 0.014) (**Suppl. Fig. 5E**), while Tb.N_CONT_ showed a strong positive alignment with Lac.Circ_CONT_ (*r* = 0.675, *p* = 0.016) (**Suppl. Fig. 5F**). Conversely, Host genome copy_CONT_ displayed a statistically significant negative correlation with Tb.BMD_CONT_ (*r* = −0.633, *p* = 0.027) (**Suppl. Fig. 5G**).

Within the cortical bone compartment during physiological homeostasis (**Suppl. Fig. 6A**), %DBM_CONT_ exhibited significant negative correlations with Ct.BV/TV_CONT_ (*r* = −0.701, *p* = 0.011) (**Suppl. Fig. 6B**), Ct.BMD_CONT_ (*r* = −0.756, *p* = 0.004) (**Suppl. Fig. 6C**), Ct.Ps.BV/TV_CONT_ (*r* = −0.694, *p* = 0.012) (**Suppl. Fig. 6D)**, and Ct.Ps.BMD_CONT_ (*r* = −0.794, *p* = 0.002) (**Suppl. Fig. 6E**). Correspondingly, %DBM_CONT_ displayed positive linear associations with Ct.Po_CONT_ (*r* = 0.701, *p* = 0.011) (**Suppl. Fig. 6F**) and Ct.Ps.Po_CONT_ (*r* = 0.694, *p* = 0.012) (**Suppl. Fig. 6G**). Additional significant linear inter-relationships among homeostatic cortical bone microarchitectural parameters were identified in the correlation matrix (**Suppl. Fig. 6A**), with key representative regressions delineated in **Suppl. Figure 6B-6G**.

### Relationships in trabecular bone between microbial load, structure and host tissue response during infection

To investigate the relationship between bone destruction, microbial load, and subsequent host cellular and morphometric responses during infection, comprehensive Pearson correlation and linear regression analyses were performed across multi-scale datasets. Matrices illustrating the overall Pearson correlation coefficients (*r*) across all micro-CT, histological, and molecular parameters for trabecular bone in the infection group are presented in **Fig. 5A**. Positive linear relationships were maintained between Tb.BV/TV and Tb.Th in both infection (*r*=0.677, *p*=0.016) and control (*r*=0.645, *p*=0.023) group (**Fig. 5B**), as well as between Tb.BV/TV and Tb.BMD (INF: *r*=0.944, *p*<0.0001; CONT: *r*=0.885, *p*=0.0001) (**Fig. 5C**) and between Tb.Th and Tb.BMD (INF: *r*=0.799, *p*=0.0002; CONT: *r*=0.682, *p*=0.014) **(Fig. 5D**). However, Tb.N correlated positively with Tb.BMD (*r*=0.589, *p*=0.039) (**Fig. 5E**) and Tb.Th (*r*=0.7336, *p*=0.0006) (**Fig. 5F**) in the infected mice, whereas these relationships were absent in control mice. %DBM also showed a significant positive correlation with Tb.Th (*r*=0.7717, *p*=0.009) in the infection group but not in controls (*r*=−0.088, *p*=0.785) (**Fig. 5G**). Bacterial genome copy number exhibited a weak negative linear correlation with Tb,BV/TV (*r*=−0.576, *p*=0.050) in the infection group (**Fig. 5H**).

**Figure 5.**
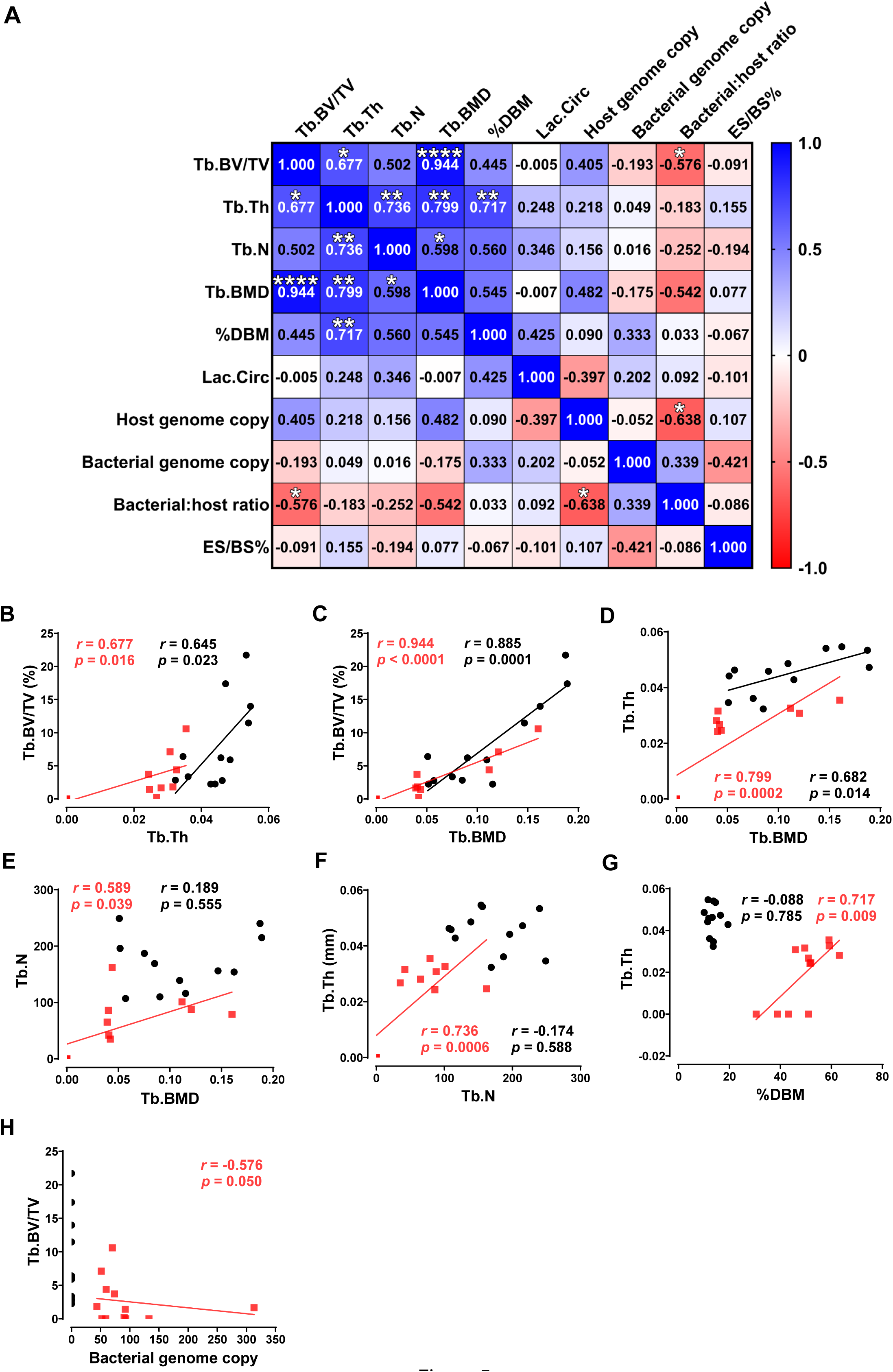
Multi-parameter correlation and regression analysis between bacterial load, trabecular microarchitecture, and histological parameters in the ipsilateral tibia of the infection group. (A) Pearson correlation heatmap illustrating the relationships among trabecular bone micro-CT parameters (Tb.BV/TV, Tb.Th, Tb.N, Tb.BMD), histological metrics (%DBM, Lac.Circ, ES/BS%), and molecular metrics (Host genome copy, Bacterial genome copy, Bacterial:host ratio) in the infected ipsilateral tibia (n=12, pooled from 6 males and 6 females). The colour scale represents the Pearson correlation (*r*), ranging from red (strong negative correlation, −1.0) to blue (strong positive correlation, 1.0). Asterisks statistical significance: \**p* < 0.05, \*\**p* < 0.01, \*\*\**p* < 0.001, \*\*\*\**p* < 0.0001. (B-H) Representative linear regression scatter plots comparing structural and matrix relationships between infected mice (red squares) and control mice (black circles). Fitted regression lines are displayed for relationships exhibiting statistically significant correlations (*p*<0.05), whereas scatter points without regression lines indicate non-significant alignments (*p*>0.05). Pearson correlation coefficient (*r*) and the significance level (*p-value*) are colour-coded for infection (red) and control (black) groups in each panel.

### Relationships in cortical bone between microbial load, structure and host tissue response during infection

Matrices illustrating the overall Pearson correlation coefficients (*r*) across all micro-CT, histological, and molecular parameters for cortical bone in the infection group are presented in **Fig. 6A**. In the cortical compartment, host cell genome copy exhibited statistically significant positive linear correlations with Ct.BMD (*r*=0.623, *p*=0.030) (**Fig.6B**), Ct.Ps.BV/TV (*r*=0.593, *p*=0.042) (**Fig.6C**), and Ct.Ps.BMD (*r*=0.694, *p*=0.0012) (**Fig.6D**), alongside a significant negative linear correlation with Ct.Ps.Po (*r*=−0.593, *p*=0.042) (**Fig.6E**) in the infection group. In contrast, the control group showed no significant linear associations across these corresponding host cellularity plots.

**Figure 6.**
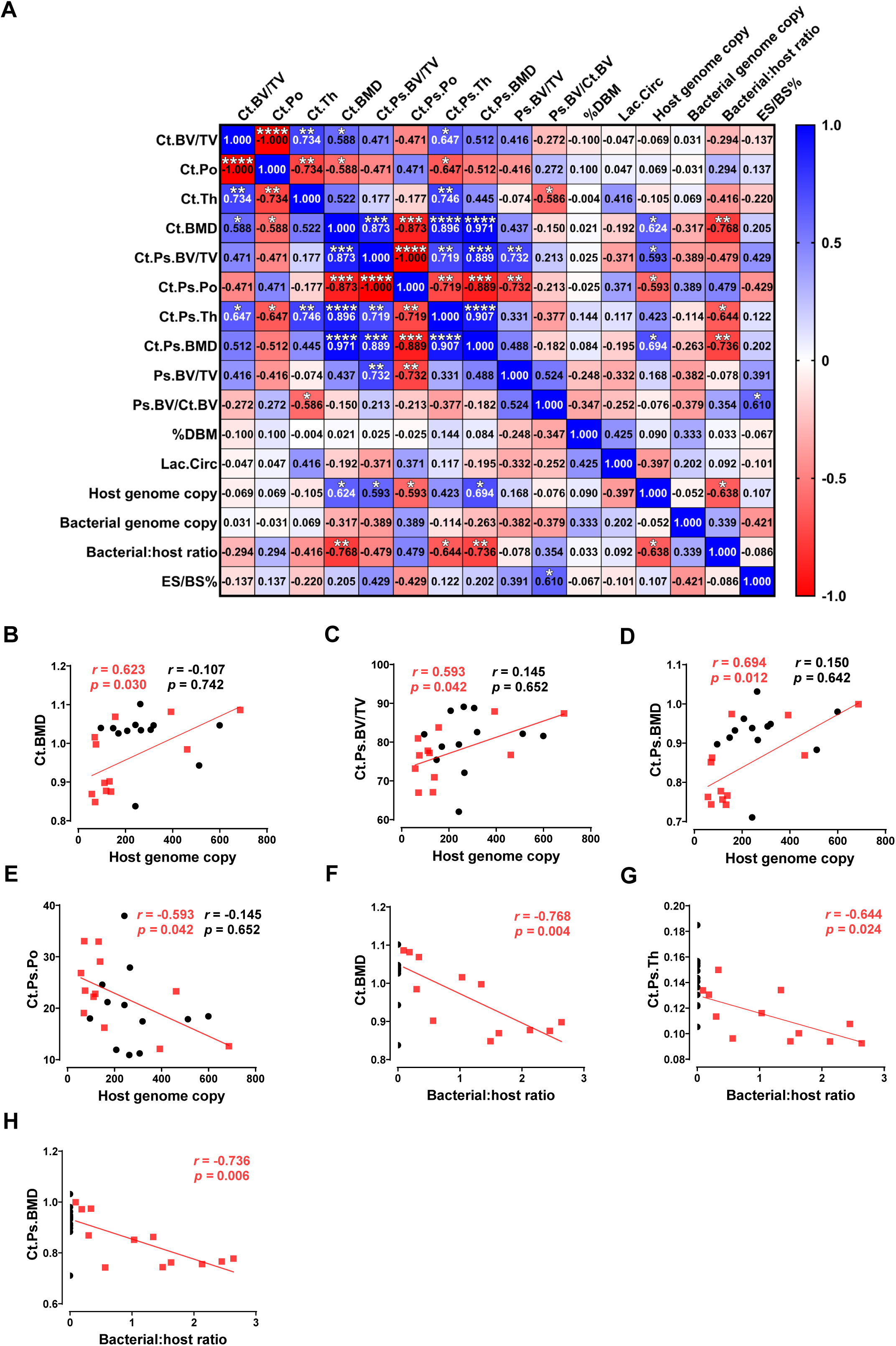
Multi-parameter correlation and linear regression analyses among cortical bone microarchitecture, histological metrics, and molecular load in the ipsilateral tibia of the infection group. (A) Pearson correlation heatmap showing the relationships among cortical micro-CT parameters (Ct parameters, Ct.Ps parameters, Ps parameters), histological metrics (%DBM, Lac.Circ, ES/BS%), and molecular metrics (Host genome copy, Bacterial genome copy, Bacterial:host ratio) in the infected ipsilateral tibia (n=12 pooled from 6 males and 6 females). The colour scale indicates the Pearson correlation coefficient (*r*), ranging from red (strong negative correlation, −1.0) to blue (strong positive correlation, 1.0). Asterisks indicate statistical significance for the correlations (\**p*<0.05, \*\**p*<0.01, \*\*\**p*<0.001, \*\*\*\**p*<0.0001). (B-H) Representative linear regression scatter plots highlighting structural, histological, and pathogen-load relationships. Data points and fitted regression lines are presented for the infection group (red squares and red lines) alongside controls (black circles; black lines are shown only where significant linear correlations exist, *p*<0.05). Pearson correlation coefficient (*r*) and the significance level (*p-value*) for both infection (red) and control (black) groups are provided in each panel.

There were no observed relationships between host measures and unadjusted bacterial genome copy number. However, the bacterial:host genome ratio in the infection group displayed strong, statistically significant negative linear regressions with Ct.BMD (*r*=−0.768, *p*=0.004) (**Fig. 6F**), Ct.Ps.Th (*r*=−0.644, *p*=0.024) (**Fig.6G**), and Ct.Ps.BMD (*r*=−0.736, *p*=0.006) (**Fig. 6H**), consistent with the host cell response to infection being the predominant driver of histopathological changes over pathogen load.

## Discussion

This study presents a quantified evaluation framework of implant-associated bone infection that integrates automated micro-CT segmentation for quantifying changes to cortical, trabecular and periosteal reactive bone compartments, digitalised RGB histopathology for assessing effects on host cells and the bone matrix, and molecular quantification of pathogen inoculum and tissue load using ddPCR. Importantly, our analyses challenge the traditional binary view of pathological bone remodelling governed exclusively by osteoclasts and osteoblasts.

The precise segmentation of cortical and trabecular bone is fundamental to the radiographic analysis of bone infection. Traditional manual tracing is not only time-consuming but also prone to observer bias [22], particularly at the endocortical interface, where infection-induced osteolysis and sequestra may blur anatomical boundaries [23]. Here, we established a standardised, automated task-list that enables the objective isolation of cortical, trabecular, and periosteal reactive bone compartments. This protocol significantly reduces processing time and minimises subjective interference, ensuring a more accurate and reproducible compartmentalised analysis than conventional manual contouring. The quantitative micro-CT findings revealed that trabecular bone experienced more pronounced resorption with the progression of infection time. In contrast, within the cortical compartment, a dynamic compensatory remodelling pattern was observed. While the Ct.BV/TV declined and Ct.Po increased significantly over time, indicating progressive internal structural destruction, the Ct.Ps captured a transient dip at day 7 followed by persistent periosteal adaptation. This phenotypic buffering of the overall cortical envelope is primarily attributed to the continuous accumulation of reactive periosteal bone (Ps.BV/Ct.BV) over the course of infection, which appears to compensate for the severe loss of the original cortex, providing a detailed quantitative signature of structural adaptation during implant-associated bone infection.

Beyond micro-CT alterations, we have reported that bone matrix degradation and the reaction within the osteocyte lacunocanalicular microenvironment are hallmark features of human PJI [2, 3, 21]. Our recent clinical findings demonstrated that %DBM, osteocyte lacunar morphology and canalicular measures serve as histopathological indicators to differentiate human PJI from aseptic osteolysis, with PJI patients exhibiting significantly more extensive collagen breakdown and increased osteocyte lacunar circularity, among several other measures [3]. The novel pathological features observed in the infection group of this mouse model are highly consistent with these clinical observations. Utilising digitalised RGB staining, we demonstrated marked increases in %DBM, Lac.Area, Lac.Perim, and Lac.Circ, accompanied by a significant reduction in Lac.Asp.Ratio under infection conditions. Mechanistically, these tissue-level phenotypes align tightly with our previous findings in human osteocytes and bone tissue that *S. aureus* exposure strongly upregulates the expression of matrix metalloproteinases (MMP-1, MMP-3 and MMP-13) and cathepsin K in osteocytes; the localised secretion of these mediators drives active bone matrix breakdown in a form of pathological osteocytic osteolysis [2]. This targeted enzymatic erosion appears also to effectively remodel the original lacunar boundaries, shifting the lacunar geometry toward a circular morphology (Lac.Circ) within the degraded or poorly organised bone matrix, providing *in vivo* evidence of pathogen-driven osteocytic osteolysis. Increased osteocyte lacunar circularity (sphericity) is often associated with conditions of bone loss considered to be due to loss of mechanosensation by the osteocyte [24, 25]. We were unable to measure canaliculi using the RGB trichrome staining protocol modified for mouse bone, unlike the human equivalent [3], due to insufficient image resolution, thus it remains to be determined if these measures are also affected in the mouse model utilised here.

To bridge the critical gap between direct microbial colonisation and pathobiological skeletal destruction, ddPCR-based quantitative molecular profiling was integrated with multi-compartmental structural and histological metrics. A key advantage of ddPCR over traditional culture-based colony-forming unit (CFU) assays lies in its ability to quantify total genomic load independent of bacterial culturability, mitigating false negatives caused by intracellular persistence or viable-but-non-culturable (VBNC) small-colony variants (SCVs) [18, 19, 21, 26]. Crucially, our multi-parameter correlation analyses revealed that neither absolute bacterial genome copies nor classical osteoclast activity (ES/BS%) in isolation fully aligned with the magnitude of structural decay. Instead, under infection conditions, the physiological homeostatic coupling evident in uninfected control bone was fundamentally disrupted and reconfigured.

In the healthy control compartment, bone microarchitecture obeyed tightly regulated physiological alignments, for example, strong positive coupling observed between Tb.BV/TV, Tb.Th and Tb.BMD (see **Suppl. Fig. 5B–D**). Upon infection, however, the bacteria-to-host genome ratio emerged as the dominant driver of cortical destruction, exhibiting robust negative linear regressions with Ct.BMD, Ct.Ps.Th and Ct.Ps.BMD (**Fig. 6F–H**). Rather than adhering strictly to a classical two-dimensional ‘osteoblast-anabolic versus osteoclast-catabolic’ axis, these quantitative alignments underscore that infectious bone destruction is a complex, multifactorial process where traditional coupled remodelling is overlaid by pathogen-induced host cellular responses. Indeed, they highlight the pivotal immunometabolic role of the osteoblast-osteocyte lineage acting as an integrated stromal immune network [27]. Upon sensing pathogen-associated molecular patterns (PAMPs) via pattern recognition receptors, resident bone cells transition into active immune effectors, triggering local pro-inflammatory chemokine/cytokine cascades and matrix-degrading enzymes that drive localised osteolysis [2, 27]. Central to this localised catabolic response is the direct involvement of osteocytic osteolysis in dictating matrix integrity and compartment remodelling. As the most abundant resident cell population within the mineralised matrix, osteocytes do not merely serve passive mechanosensory functions; under *S. aureus* exposure, they undergo profound morphological distortion toward circularity (Lac.Circ) and lacunar enlargement (Lac.Area), accompanied by increased %DBM (see **Fig. 4**). In infected trabecular bone, %DBM shifted from a baseline disassociation to a strong positive alignment with Tb.Th (see **Fig. 5G**). Mechanistically, this reflects active perilacunar enzymatic digestion of the surrounding bone matrix driven by osteocytic upregulation of matrix metalloproteinases (e.g. MMP-1, MMP-3 and MMP-13), as well as Cathepsin K [2, 3]. This localised, rapid perilacunar matrix breakdown precedes, and likely complements, slower-acting osteoclastic bone resorption, resulting in a rapid increase in cortical porosity (Ct.Po). Taken together, these multi-scale correlations demonstrate that osteocytic osteolysis and host stromal immune responses, rather than absolute microbial mass alone, serve as the central mechanistic engine orchestrating pathological bone destruction and reactive modelling during implant-related bone infection.

Despite the methodological advancements and mechanistic insights presented in this study, several limitations should be acknowledged. The use of 8-week-old mice during a period of rapid skeletal growth is a potential confounding factor, as intense physiological bone remodelling would overlap with infection-induced pathological remodelling. Additionally, this study utilised a fixed initial bacterial inoculum with a single reference strain, and histological/molecular evaluations were restricted to the terminal day 12 endpoint, limiting the assessment of multi-stage immune continuum dynamics. Nevertheless, these limitations were partially mitigated by leveraging longitudinal *in vivo* micro-CT within the same animals, enabling high-precision paired evaluation.

In conclusion, this study established and validated a fully quantified evaluation framework that successfully transitions the characterisation of implant-associated bone infection from qualitative description to precise, compartmentalised quantification. Multi-parameter correlation and regression analyses demonstrated that the normalised bacterial:host genome ratio, rather than absolute pathogen load, is significantly associated with reductions in cortical and periosteal BMD, while osteocyte lacunae exhibit marked pathological remodelling toward circularity (%DBM and Lac.Circ). These findings provide compelling *in vivo* evidence that, alongside classical osteoclast- and osteoblast-mediated coupled remodelling, resident osteocytes actively participate in and contribute to infection-induced bone pathology through localised matrix degradation and cellular microenvironment remodelling. This integrated platform represents a scalable methodological toolkit for unravelling complex pathobiological mechanisms in deep bone infections and evaluating novel targeted therapeutic strategies.

## Acknowledgements

The authors would like to thank the Core Animal Facility (CAF) of Adelaide University for providing animal housing and Micro-CT scanning services. We are grateful to the Adelaide Microscopy AHMS facility for providing the image analysis software and computational resources for Micro-CT data processing. Special thanks are also extended to Mr. Adnan Mulaibrahimovic (Research Technician, School of Medicine, College of Health) for his expert assistance in histological sectioning and specimen processing.

## Funding Statement

This work was funded by The National Health and Medical Research Council of Australia (NHMRC) Ideas Grant Scheme (ID 2011042) awarded to G.J.A. and P.H.A. Q.S. and M.A.H. were recipients of University of Adelaide Postgraduate Research Awards.

## Disclosures

G.J.A., D.M. and L.B.S. are named Inventors on PCT Patent PCT/AU2025/050524, Title: Histological Markers and Methods For Analysis of Bone Matrix Integrity. None of the other authors has any conflicts of interest, financial or otherwise, to disclose.

## Author Contributions

G.J.A. and P.H.A. conceived the study, contributed to experimental design and data interpretation. Q.S. principally conducted the experiments, aided by H.T. and R.S. D.M. contributed to histological methodology. Q.S. conducted statistical analysis. Q.S., M.A.H., D.M., P.H.A. and G.J.A. contributed to data visualisation. G.J.A. and P.H.A. contributed to funding acquisition. G.J.A., P.H.A. and L.B.S. supervised the study. Q.S. wrote the first draft and with G.J.A. tailored the manuscript. All authors contributed to manuscript editing and approved the final submitted version.

---

**Supplementary Table 1:**
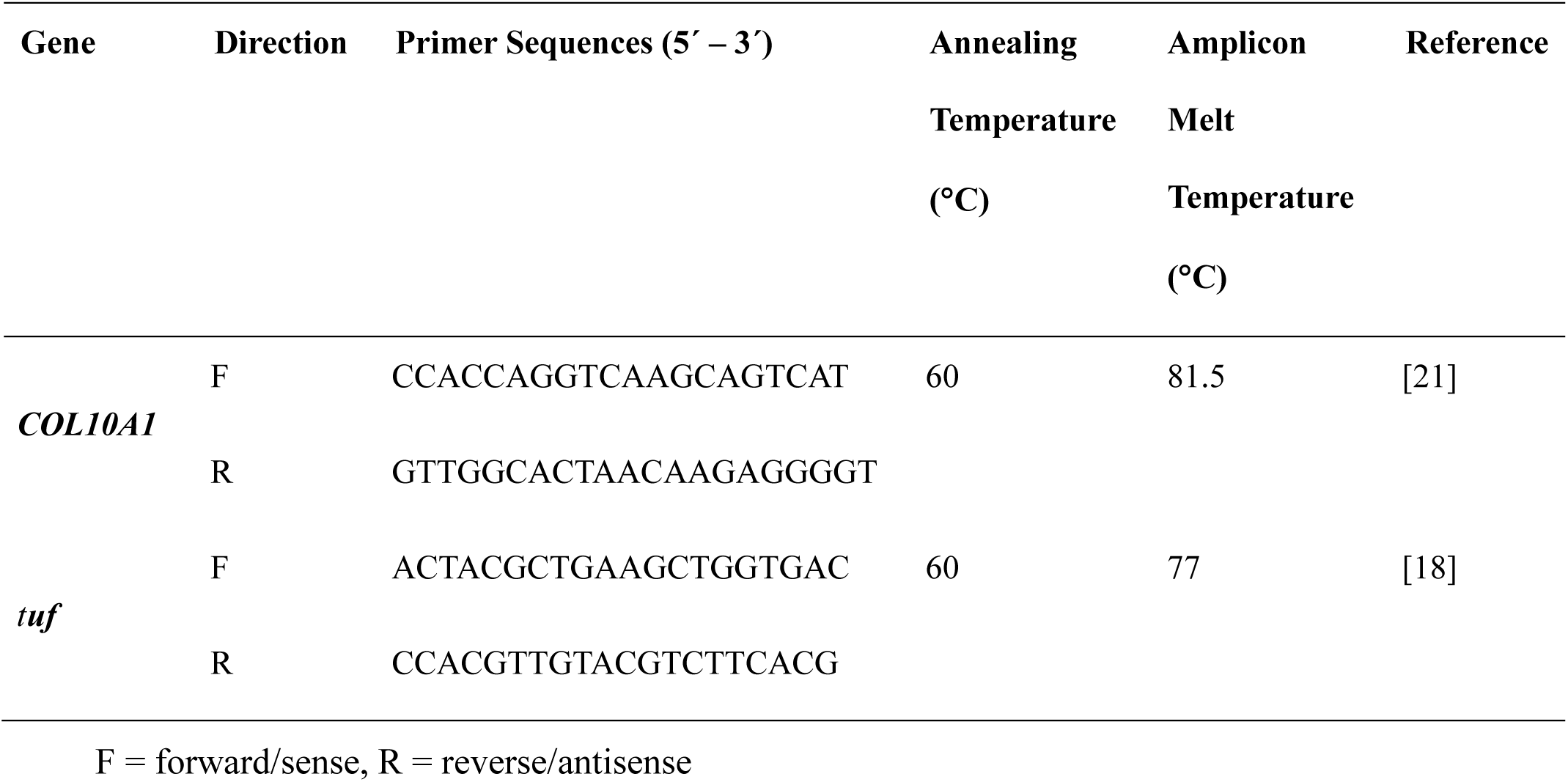
Single-copy gene DNA-specific droplet digital PCR primers used in this study.

## Supplementary Figure Legends

**Supplementary Figure 1.**
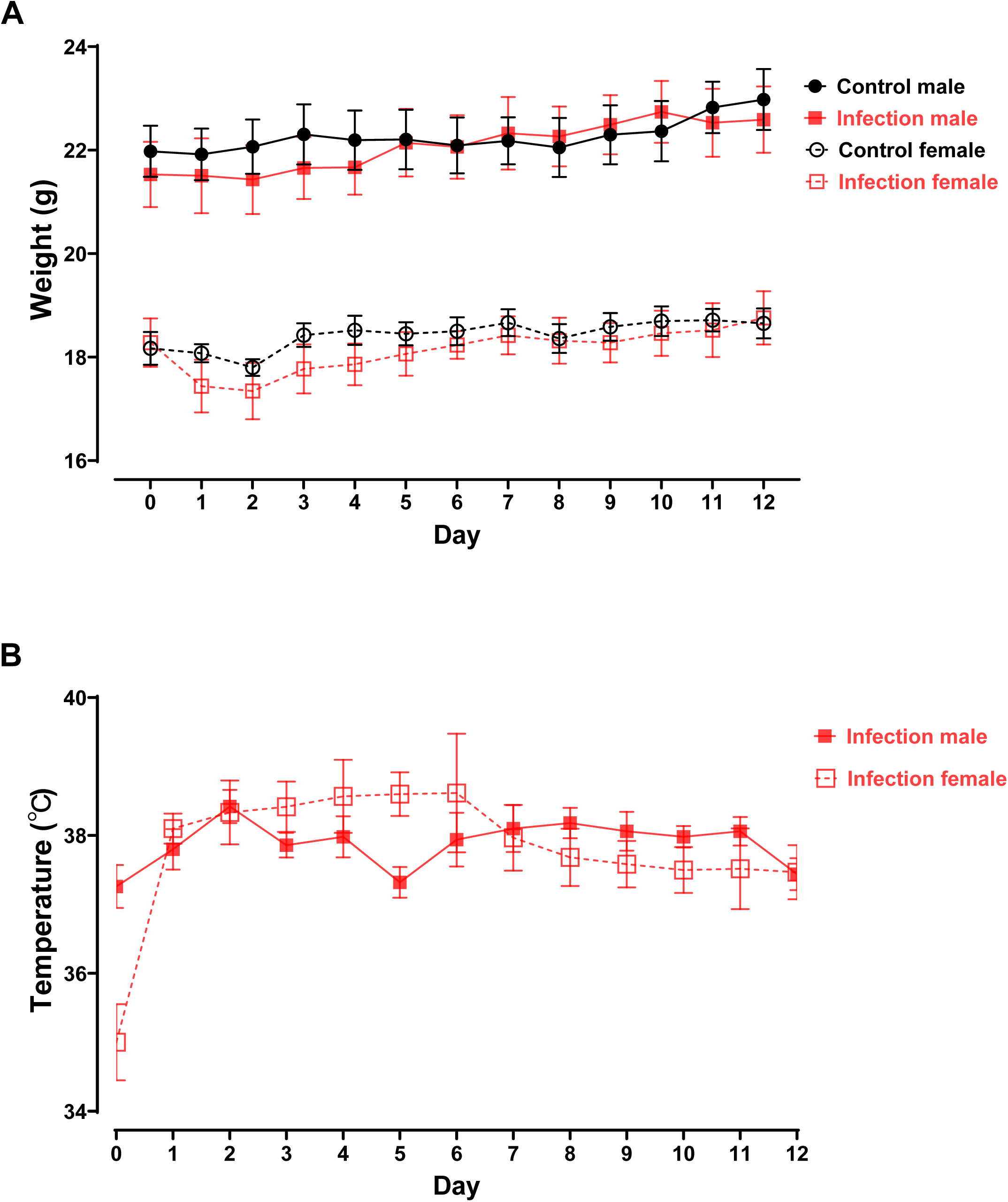
Host physiological monitoring and host genome quantification in the control group. (A) Longitudinal body weight monitoring from Day 0 to Day 12 post-surgery in male and female mice (Control male, Infection male, Control female, Infection female; n=6 per gender per group). Both control and infection groups exhibited steady body weight maintenance and recovery throughout the experimental period. (B) Postoperative body temperature profile in infected male and female mice (n=6 per gender) monitored daily from Day 0 to Day 12. All data are presented as descriptive statistics (mean ± SEM) to reflect overall host physiological stability and quantitative baseline values; no formal inferential hypothesis testing was performed.

**Supplementary Figure 2.**
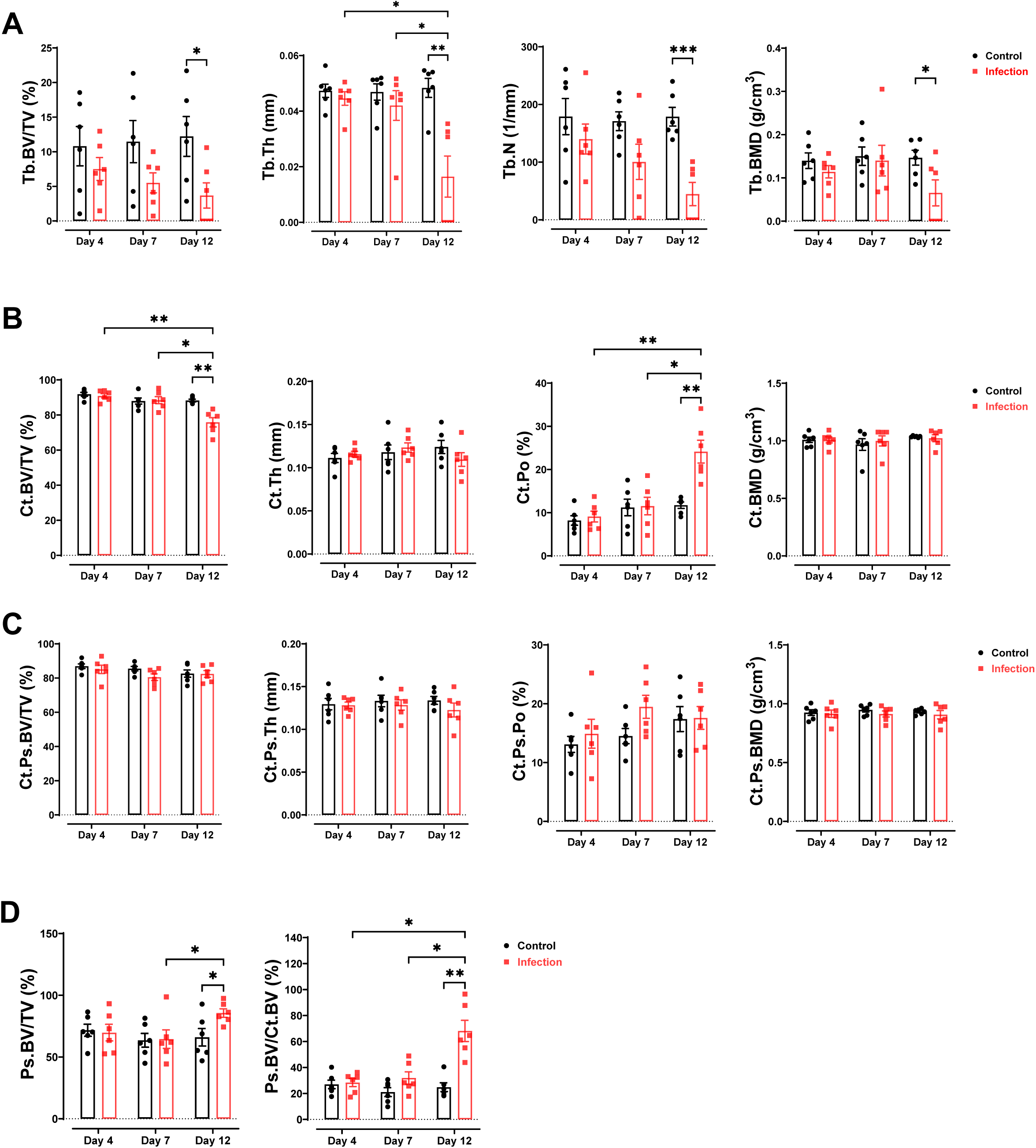
Longitudinal micro-CT evaluation of trabecular and cortical bone alterations in male mice. (A) Quantitative analysis of trabecular bone parameters (Tb.BV/TV, Tb.Th, Tb.N, and Tb.BMD) in male control and infection groups at post-operative days 4, 7, and 12 (n=6 male mice per group). (B) Quantitative micro-CT evaluation of the cortical bone compartment excluding periosteal reactive bone (Ct.BV/TV, Ct.Th, Ct.Po, and Ct.BMD). (C) Quantitative analysis of combined cortical and periosteal reactive bone (Ct.Ps.BV/TV, Ct.Ps.Th, Ct.Ps.Po, and Ct.Ps.BMD). (D) Assessment of reactive periosteal bone formation (Ps.BV/TV and Ps.BV/Ct.BV). Data are presented as mean ± SEM (n=6 male mice per group) to complement the pooled datasets shown in Figures 2 and 3. Statistical significance was determined by Two-way ANOVA followed by Tukey’s multiple comparisons test. \**p*<0.05, \*\**p*<0.01, \*\*\**p*<0.001 indicate significant differences between the indicated groups or time points.

**Supplementary Figure 3.**
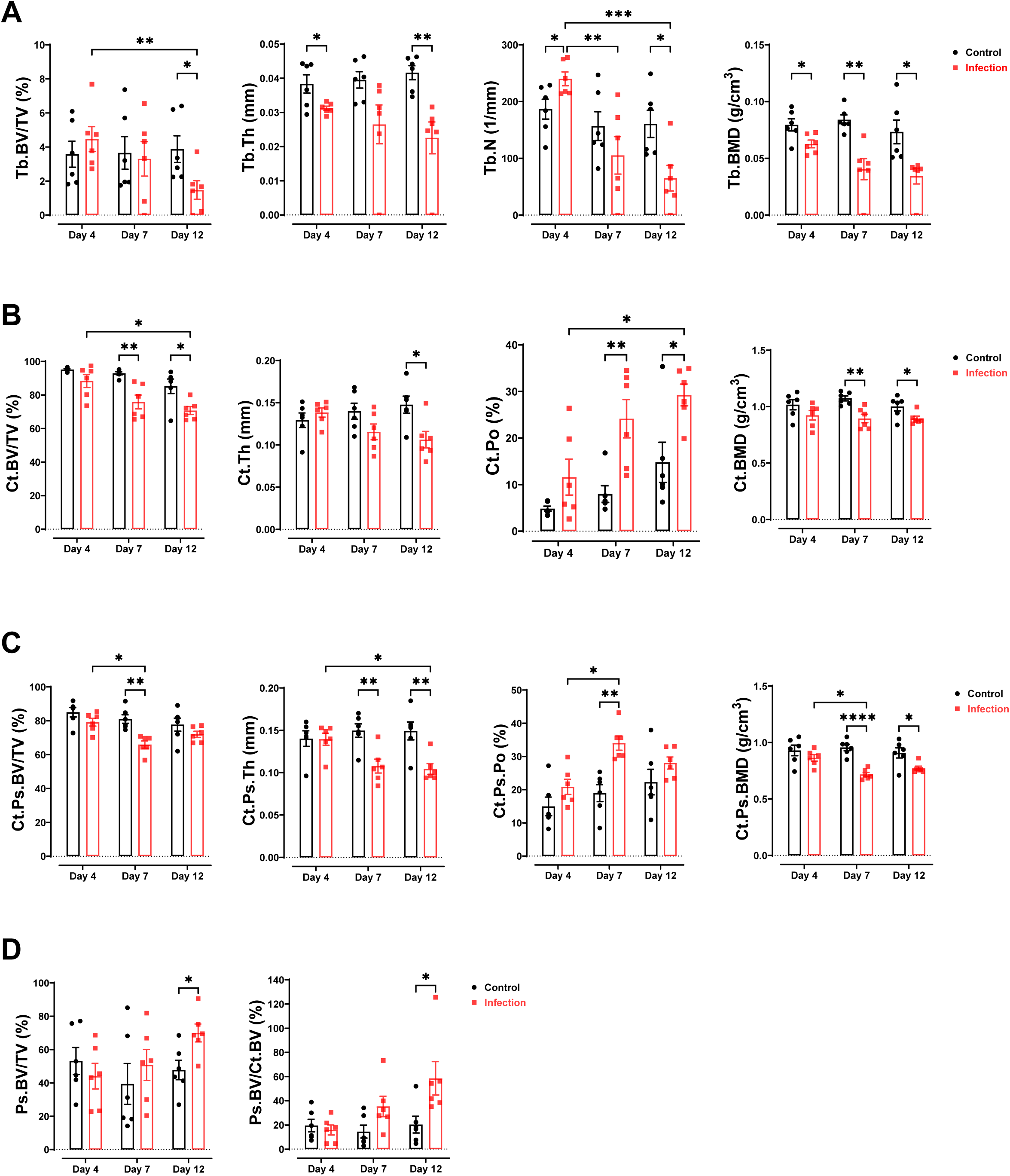
Longitudinal micro-CT evaluation of trabecular and cortical bone alterations in female mice. **(A)** Quantitative analysis of trabecular bone parameters (Tb.BV/TV, Tb.Th, Tb.N, and Tb.BMD) in female control and infection groups at post-operative days 4, 7, and 12 (n=6 female mice per group). **(B)** Quantitative micro-CT evaluation of the cortical bone compartment excluding periosteal reactive bone (Ct.BV/TV, Ct.Th, Ct.Po, and Ct.BMD). **(C)** Quantitative analysis of combined cortical and periosteal reactive bone (Ct.Ps.BV/TV, Ct.Ps.Th, Ct.Ps.Po, and Ct.Ps.BMD). **(D)** Assessment of reactive periosteal bone formation (Ps.BV/TV and Ps.BV/Ct.BV). Data are presented as mean ± SEM (n=6 female mice per group) to complement the pooled datasets shown in Figures 2 and 3. Statistical significance was determined by Two-way ANOVA followed by Tukey’s multiple comparisons test. *p*<0.05, \*\**p*<0.01, \*\*\**p*<0.001, \*\*\*\**p*<0.0001 indicate significant differences between the indicated groups or time points.

**Supplementary Figure 4.**
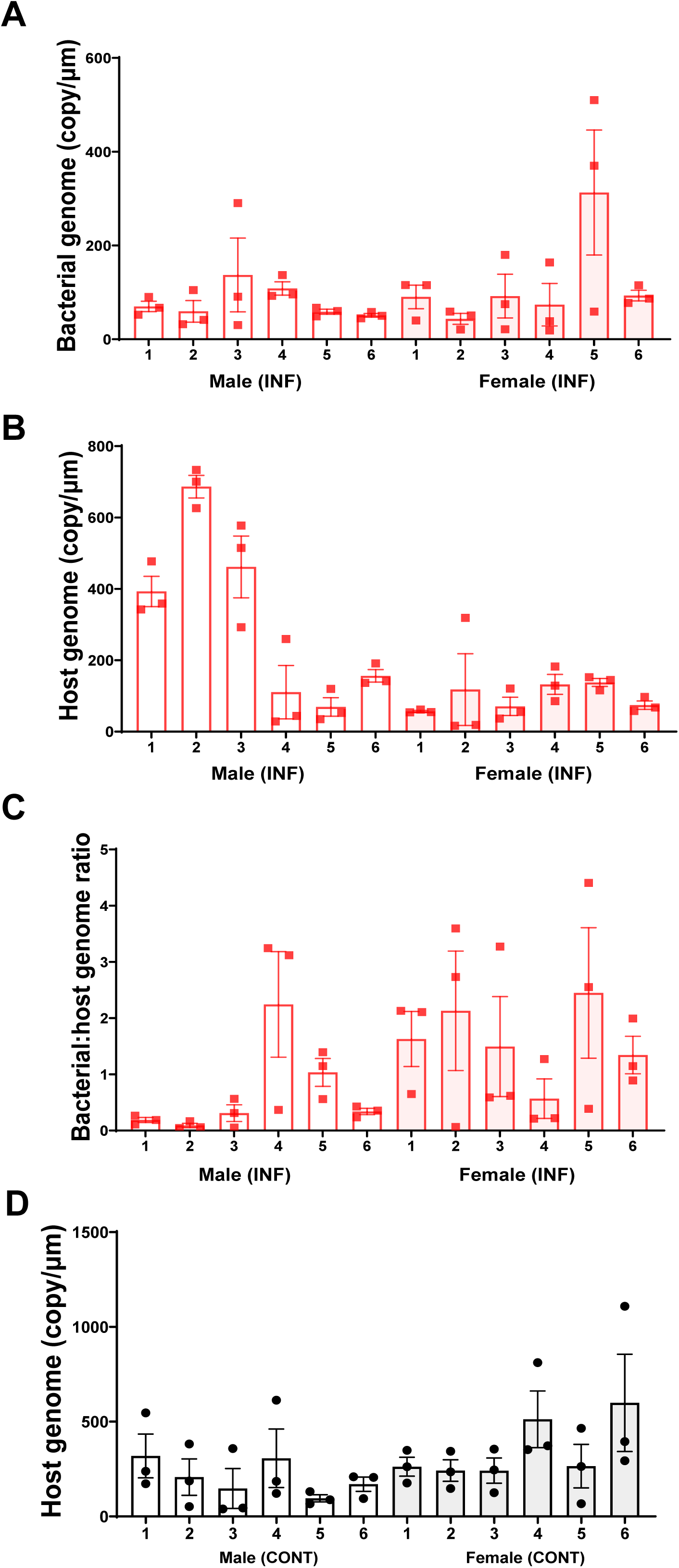
Molecular quantification of bacterial and host genome copy (n = 12, comprising 6 males and 6 females). (A) Quantification of bacterial genome copies and (B) host genome copies performed using ddPCR on DNA extracted from three independent pathological sections per mouse (section thickness = 20µm) in the infection group. (C) The bacterial load expressed as the ratio of bacterial to host genome copies to account for inter-individual and intra-tissue variability in the infection group. (D) Quantification of host genome copies via ddPCR in histological sections from control mice (n=12, comprising 6 males and 6 females) in the control group. Data are presented as mean ± SEM for each mouse, with three data points representing the independent section levels. Statistical significance of the bacterial-to-host genome ratio among individual mice was assessed using one-way ANOVA followed by Tukey’s multiple comparisons test (no statistically significant differences were detected between individuals).

**Supplementary Figure 5.**
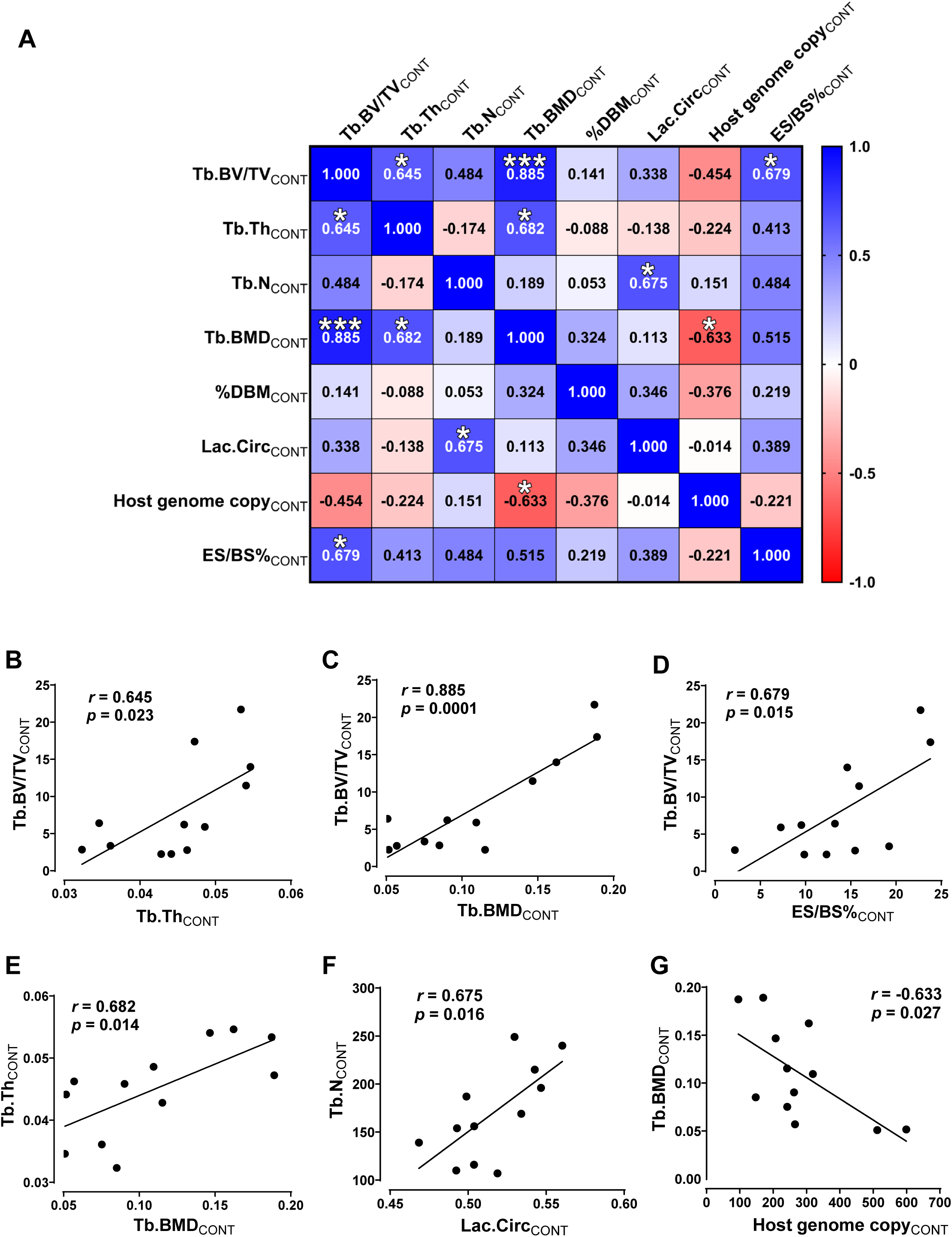
Multi-parameter correlation and linear regression analyses among trabecular bone microarchitecture, histological metrics and host cellularity in ipsilateral tibia of the control group. (A) Pearson correlation heatmap illustrating physiological relationships among trabecular micro-CT parameters (Tb.BV/TV_CONT_, Tb.Th_CONT_, Tb.N_CONT_, Tb.BMD_CONT_), histological metrics (%DBM_CONT_, Lac.Circ_CONT_, ES/BS%_CONT_), and molecular metrics (Host genome copy_CONT_) in the control group. The colour scale indicates the Pearson correlation coefficient (*r*), ranging from red (strong negative correlation, −1.0) to blue (strong positive correlation, +1.0). Asterisks indicate statistical significance (\**p*<0.05, \*\*\**p*<0.001). (B) Linear regression plots displaying statistically significant correlations under homeostatic conditions. Individual data points with fitted regression lines, Pearson correlation coefficients (*r*), and *p*-values are shown in each plot.

**Supplementary Figure 6.**
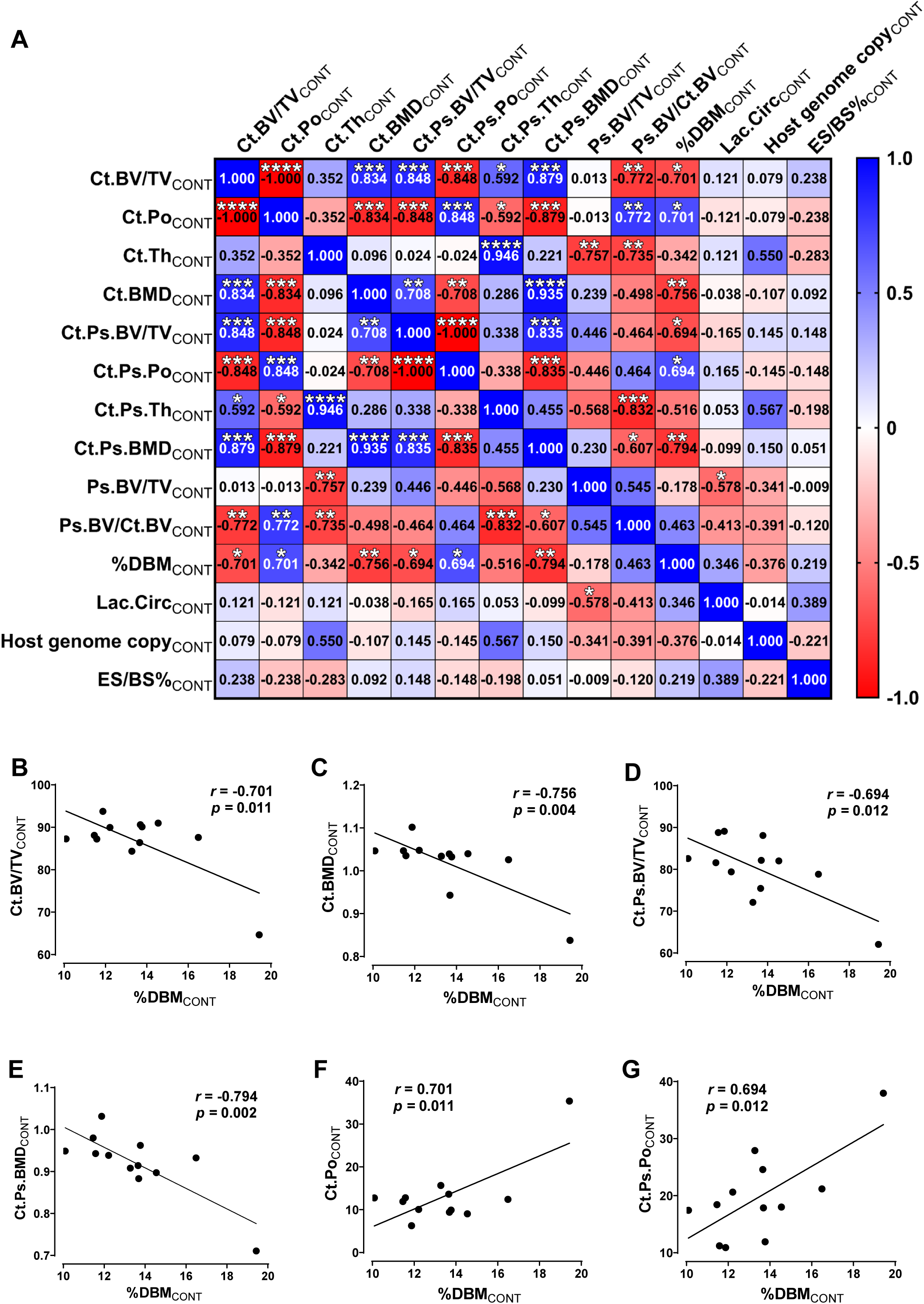
Multi-parameter correlation and linear regression analyses among cortical bone microarchitecture, histological metrics, and host cellularity in the ipsilateral tibia of the control group. (A) Pearson correlation heatmap illustrating physiological relationships among cortical micro-CT parameters (Ct_CONT_ parameters, Ct.Ps_CONT_ parameters, and Ps_CONT_ parameters), histological measures (%DBM_CONT_, Lac.Circ_CONT_, ES/BS%_CONT_), and host molecular load (Host genome copy_CONT_) in the control group. The colour scale indicates the Pearson correlation coefficient (*r*), ranging from red (strong negative correlation, −1.0) to blue (strong positive correlation, +1.0). Asterisks indicate statistical significance (\**p*<0.05, \*\**p*<0.01, \*\*\**p*<0.001, \*\*\*\**p*<0.0001). (B) Representative linear regression plots displaying statistically significant homeostatic correlations in cortical bone. Individual data points with fitted regression lines, Pearson correlation coefficients (*r*), and *p*-values are shown in each plot.

